# Proteomic characterization of spinal cord synaptoneurosomes from Tg-SOD1/G93A mice supports a role for MNK1 and local translation in the early stages of amyotrophic lateral sclerosis

**DOI:** 10.1101/2022.03.30.486360

**Authors:** Juan José Casañas, María Luz Montesinos

**Affiliations:** Departamento de Fisiología Médica y Biofísica, Universidad de Sevilla, E-41009 Sevilla, Spain; Instituto de Biomedicina de Sevilla, IBIS/Hospital Universitario Virgen del Rocío/CSIC/Universidad de Sevilla, Sevilla, Spain

**Keywords:** ALS, iTRAQ, local translation, MNK1, SOD1, spinal cord

## Abstract

The isolation of synaptoneurosomes (SNs) represents a useful means to study synaptic events. However, the size and density of synapses varies in different regions of the central nervous system (CNS), and this also depends on the experimental species studied, making it difficult to define a generic protocol for SNs preparation. To characterize synaptic failure in the spinal cord (SC) in the Tg-SOD1/G93A mouse model of amyotrophic lateral sclerosis (ALS), we applied a method we originally designed to isolate cortical and hippocampal SNs to SC tissue. Interestingly, we found that the SC SNs were isolated in a different gradient fraction to the cortical/hippocampal SNs. We compared the relative levels of synaptoneurosomal proteins in control (Tg-SOD1) animals, with Tg-SOD1/G93A mice at onset and those that were symptomatic using iTRAQ proteomics. The results obtained suggest that an important regulator of local synaptic translation, MNK1 (MAP kinase interacting serine/threonine kinase 1), might well influence the early stages of ALS.

## INTRODUCTION

Amyotrophic Lateral Sclerosis (ALS) is a fatal disease that affects both upper and lower motor neurons, producing muscular atrophy, paralysis and finally, death due to respiratory failure within 5 years of the first disease symptoms appearing. Although in most cases the origin of ALS is unknown, 10% of ALS cases are familial and of these, 20-25% are due to dominant mutations affecting the SOD1 (Superoxide Dismutase 1) gene [1]. Moreover, SOD1 has also been implicated in 1-7% of the sporadic forms of ALS.

Neuronal degeneration is not apparent in SOD1 knockout mice [2] and many of the mutant SOD1 proteins that do produce neurodegeneration exhibit normal enzymatic activity [3, 4]. Indeed, the neurotoxicity of these mutant proteins seems to be related to the acquisition of a new property, the aggregation and formation of mutant SOD1 protein inclusions [5]. These protein aggregates are in line with the aggregates associated with other neurodegenerative diseases like Alzheimer’s, Huntington’s and Parkinson’s disease. The tendency to form SOD1 aggregates seems to be restricted to motor neurons, as they do not appear in hippocampal or dorsal root ganglion (DRG) neurons transfected with mutant SOD1 constructs, only in transfected motor neurons [6]. A number of hypotheses have been proposed to explain the toxicity of SOD1 aggregates in motor neurons, including: the co-aggregation of essential cytoplasmic proteins; the clogging of the proteasome with misfolded proteins; a depletion of chaperones; the dysfunction of mitochondria and other cellular organelles (for a review see [7]). However, it remains unclear if these phenomena are a cause or a consequence of the disease.

Among the various models of ALS developed, the Tg-SOD1/G93A [8] transgenic mouse model is probably one of the best characterized. This transgenic line carries a mutant version of the human SOD1 gene in which the glycine at position 93 is replaced by alanine. Interestingly, dendrites with aggregates of the mutant SOD protein can initially be seen in Tg-SOD1/G93A mice at pre-symptomatic stages [9], as also seen in other murine models of ALS [10, 11]. Significantly, dendritic aggregates have also been observed in tissue from individuals diagnosed with ALS [12, 13]. We previously found hippocampal synaptoneurosomes (SNs) to be strongly enriched in SOD1 mRNA, suggesting that it could be translated locally in dendrites (unpublished data). Accordingly, FMRP (Fragile-X mental retardation protein), a key local regulator of translation [14], has been shown to activate SOD1 translation [15]. Finally, SOD1 mutations in the 3’UTR (untranslated region) have been linked to ALS [16, 17]. It is known that motifs in the 3’UTR regulate the transport, stability and local translation of dendritic mRNAs (reviewed in [18]). In fact, evidence is accumulating that supports a role for RNA metabolism in ALS, not least since many genes mutated in ALS are RNA-binding proteins, including TDP-43 [19], FUS [20, 21], ATXN2 [22], hnRNPA1 and hnRNPA2/B1 [23]. Indeed, the inclusions from ALS patients contain RNA and RNA-binding proteins (reviewed in [24]). Together, these observations suggest that altered local dendritic mRNA translation could be one of the initial events involved in ALS motor neuron degeneration.

As a first step to investigate this hypothesis, we used here an improved protocol to obtain spinal cord (SC) SNs that we had originally designed to isolate cortical and hippocampal SNs [25]. Remarkably, SC SNs were located in a different fraction to that of the hippocampal SNs, which we characterized both biochemically and by electron microscopy (EM). Using isobaric tags for relative and absolute quantitation (iTRAQ) proteomics, we compared the relative levels of proteins in SC SNs from Tg-SOD1 control mice to those from Tg-SOD1/G93A mice at the onset of their symptoms and in symptomatic mice. A significant number of the affected proteins in the samples at onset are targets of the MAP kinase interacting serine/threonine kinase 1 (MKNK1, also known as MNK1), a protein that acts in the Ras-ERK signaling pathway and that also influences FMRP-regulated mRNA translation. In fact, reduced levels of MNK1 were measured in SC SNs from Tg-SOD1/G93A mice. Finally, we found that the rates of translation in SNs from the SC decreased at the onset of the Tg-SOD1/G93A phenotype. Overall, our results suggest that deregulated local synaptic translation controlled by MNK1 might play a role in the early stages of ALS.

## MATERIALS AND METHODS

### Animals

B6SJL-Tg(SOD1)2Gur/J (Tg-SOD1) and B6SJL-Tg(SOD1*G93A)1Gur/J (Tg-SOD1/G93A) mice [8] were obtained from Jackson Laboratories, and they were maintained on a mixed genetic background as recommended, breeding hemizygous male carriers with B6SJLF1 females. These B6SJLF1 females were obtained by crossing C57BL/6J females with SJL/JOrlCrl males (Charles River, France). All the experiments performed here were carried out in accordance with the European Union directive (2010/63/EU) for the protection of the animals used for research purposes. All the protocols were approved by the Regional Government (Junta de Andalucía, Spain) Ethics Committee. The criteria used to classifying the Tg-SOD1/G93A animals as at disease onset or symptomatic are summarized in Table S1 (Supplemental File 1).

### SN isolation

SNs were isolated as described previously [25]. Briefly, the SC from 6-9 mice was dissected out rapidly and homogenized with a glass-teflon Dounce homogenizer in 12 ml of homogenization buffer: 10 mM Hepes [pH 7.4]; 320 mM sucrose; 1.0 mM MgSO_4_; protease inhibitors leupeptin (10 μM), aprotinin (2 μg/ml) and pepstatin (1 μM). The homogenate was centrifuged at 1,000 g for 10 min at 4 °C and the resulting pellet (P1) was resolved on an Optiprep discontinuous gradient (9-40% Optiprep). Four bands (O1 to O4: Figure 1A) were obtained after centrifugation at 16,500 g for 30 min at 4 °C. The fractions O1 and O2 were recovered and passed separately through a discontinuous Percoll gradient (10-25% Percoll). After centrifugation at 32,400 g for 20 min at 4 °C, five bands were obtained from fraction O1 and O2: 1P1 to 1P5 (Figure 1A) and 2P1 to 2P5, respectively (data not shown). Of these, fraction 1P5 corresponds to the SC SNs.

**Figure 1.**
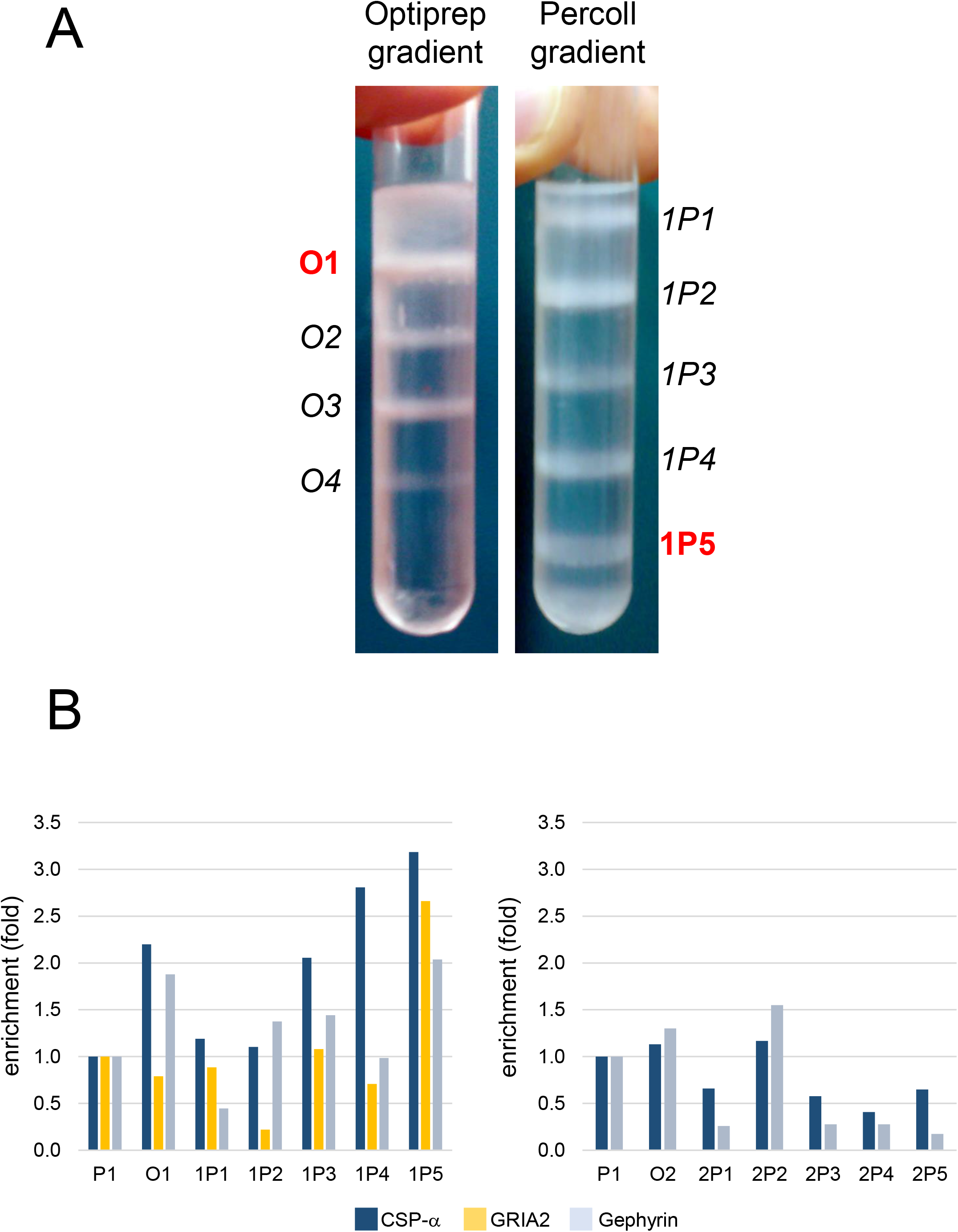
SC SNs isolation. (A) Optiprep and Percoll gradients from a typical SC SN isolation. Four subcellular fractions (O1 to O4) were obtained after the Optiprep gradient and when the material from the O1 fraction was separated on a Percoll gradient, five bands were observed (1P1 to 1P5). The SNs were found in fraction 1P5. (B) Quantification of synaptic proteins in the Percoll gradient fractions from the mouse SC. Levels of direct chemiluminescence for the CSP-α, GRIA2 and gephyrin proteins detected in P1, O1 and O2, and in the corresponding Percoll gradient fractions (1P1 to 1P5 from O1, and 2P1 to 2P5 from O2), normalized to the total protein in the samples. Normalized chemiluminescence for P1 was arbitrarily set to 1. A representative experiment is shown. Note that the GRIA2 levels for fractions O2 and 2P1 to 2P5 were not determined.

### SC protein extracts

Total lumbar SC protein extracts were prepared using a mechanical overhead stirrer (Heidolph model RZR 2020) with a 2 ml Potter-Elvehjem PTFE pestle. The tissue was homogenized in the following extraction buffer: 50 mM Tris buffer [pH 7.4], 0.5% Triton X-100, 150 mM NaCl, 1 mM EDTA, 3% SDS, protease inhibitor cocktail (1:100 dilution, P8340: Sigma), phosphatase inhibitor cocktail 2 (1:100 dilution, P5726: Sigma), and phosphatase inhibitor cocktail 3 (1:100 dilution, P0044: Sigma). The samples were then centrifuged for 15 min at 14,500 rpm using a bench-top microfuge at room temperature, and the supernatants were recovered.

### Western blotting

To characterize the SC fractions obtained through the SN isolation protocol previously described [25], B6SLJF1 mice were used. The samples from pellet P1, fractions O1 and O2, and the Percoll fractions 1P1 to 1P5, and 2P1 to 2P5) were first resolved by SDS-PAGE using Mini-PROTEAN^®^ TGX precast gels (BioRad). The proteins were then transferred to polyvinylidene difluoride (PVDF) membranes overnight (tank blotting), stained with SYPRO^®^ Ruby protein blot stain (Invitrogen), and the SYPRO Ruby signal was measured in a ChemiDoc XRS (BioRad) imager and used for normalization. In other experiments, Western blotting was carried out following a similar protocol but Mini-PROTEAN^®^ TGX Stain-Free™ precast gels (BioRad) were used. After electrophoresis, the gels were UV irradiated in a ChemiDoc XRS apparatus (BioRad) to induce the reaction of trihalo compounds in the gel with the tryptophan residues of the proteins. The resulting fluorescent signal was proportional to the total protein present in the gel, and it was measured and used for normalization.

The membranes were probed with the primary antibodies against: CSP-α (R-807, rabbit polyclonal antiserum, a gift from Dr Rafael Fernández-Chacón, Universidad de Sevilla-IBiS, Spain); GRIA2 (anti-GluA2/AMPA2#1, rabbit polyclonal antiserum, Cat. No. 182103: Synaptic Systems); gephyrin (3B11 mouse monoclonal antibody, Cat. No. 147111: Synaptic Systems); and Mnk1 (C4C1, antibody #2195: Cell Signaling Technology). The binding of HRP-conjugated secondary antibodies was revealed by ECL Plus (GE Healthcare Life Sciences) and chemiluminescence was measured using a ChemiDoc XRS apparatus (BioRad). The linearity of the chemiluminescent signals was determined in pilot experiments for each antibody using 2-fold dilutions of a protein sample. The linearity of the SYPRO Ruby signal and the trihalo/tryptophan-produced fluorescence was also assessed similarly.

### Electron microscopy (EM)

Material from the SC 1P5 fraction of B6SLJF1 mice was pelleted and fixed for 1 h at 4°C with 2% glutaraldehyde in 0.1 M cacodylate buffer (pH 7.4) for EM. The sample was then washed with cacodylate, secondarily fixed in 1% OsO_4_, gradually dehydrated and embedded in Spurr resin. Sections were acquired at 80 Kv on a ZEISS LIBRA 120 transmission electron microscope (TEM) at the CITIUS Microscopy Service (Universidad de Sevilla).

### iTRAQ labeling and analysis

SC SNs were isolated from wild type (WT) mice (littermates of Tg-SOD1/G93A animals that did not bear the transgene), from Tg-SOD1/G93A mice that had just started to show some muscle weakness (G93A-ONSET) and from overtly symptomatic animals (G93A-SYMPTOM). In order to control for any changes due to overexpression of the human SOD1 transgene, SNs were also isolated from Tg-SOD1 mice.

Protein extraction, iTRAQ labeling (using AB ScieX, Ref. 4390811) and tandem mass spectrometry (MS) analysis was carried out at the Instituto de Biomedicina de Sevilla (IBiS) Proteomics Service, as described previously [26]. Two sets of samples were included in the 8-plex experiment. In Set 1, 50 μg of the protein extract was labeled for each experimental condition, whereas Set 2 contained different amounts of protein in order to check the linearity of the procedure. Only Set 1 was considered in this work and the data were analyzed with the Proteome Discoverer 1.4 software (Thermo), setting a False Discovery Rate (FDR) for both protein and peptide identification as less than 0.01.

### Gene Ontology and Pathway Analyses

PANTHER version 10.0 [27] overrepresentation tests were performed on line (http://www.pantherdb.org), selecting the entire *Mus musculus* genome (22,320 proteins) and applying a Bonferroni correction for multiple testing. An Ingenuity Pathway Analysis (IPA) core analysis was performed for both up- and downregulated proteins, considering a 2-fold cut-off threshold. Only proteins identified with at least two unique peptides were included in the IPA, and the Ingenuity Knowledge Base (genes only) set was used as a reference, considering direct and indirect relationships. All the molecules and/or relationships analyzed were observed experimentally, either in the mouse, rat or human nervous system tissue or in cells (astrocytes or neurons). The IPA software generates networks composed of sets of up to 35 functionally-related proteins and each network is attributed a score (maximum score = 50). Only the most significant networks (score > 25) were considered here.

### Radioactive labeling of SNs

SNs were resuspended in SN buffer [28] that was supplemented with chloramphenicol (100 μg/ml) to inhibit the mitochondrial synthesis of proteins, as described previously [25, 28]. The samples were incubated for 30 min at 37 °C in the presence of the EasyTag Express Protein Labeling Mix 35S (PerkinElmer), pelleted, washed with SN buffer and resolved by SDS-PAGE on Mini-PROTEAN® TGX Stain-Free™ precast gels (BioRad). After electrophoresis, the gels were exposed to UV irradiation in a ChemiDoc XRS apparatus (BioRad) and the resulting fluorescent signal was used for normalization. The proteins were then transferred to PVDF membranes overnight, and the radiolabeled proteins were detected and quantified using a Cyclone Plus Phosphor Imager system (PerkinElmer).

### Statistical analysis

The quantitative data are presented as the mean ± SEM and the significance of the comparisons was assessed using a Student’s T-test (SigmaStat software).

## RESULTS

### SC SNs are found in an unexpected gradient fraction

We had previously reported an improved method for SN preparation [25] in which the homogenized tissue is subjected to isopycnic centrifugation on a discontinuous Optiprep gradient and some of the subcellular fractions obtained are then separated in Percoll gradients. When the mouse hippocampus or neocortex is used as the starting material, the SN enriched fraction corresponds to the 1P4 fraction [25], that is the fourth Percoll gradient band obtained from processing the first Optiprep gradient fraction (O1). To test whether this protocol was suitable to isolate SNs from the SC, the subcellular fractions O1, O2, 1P1 to 1P5 (Percoll gradient of O1) and 2P1 to 2P5 (Percoll gradient of O2) obtained from the SCs of adult mice (Figure 1A) were characterized in Western blots. CSP-α was used as pre-synaptic marker, whereas GluA2 and gephyrin were used as excitatory and inhibitory post-synaptic markers, respectively. Somewhat surprisingly, the strongest enrichment of pre- and post-synaptic proteins was evident not in the 1P4 but in the 1P5 fraction (Figure 1B). Indeed, TEM analysis of the material in the SC 1P5 fraction showed it was enriched in SNs. Notably, in these preparations the pre- and post-synaptic elements remained attached, and synaptic vesicles and mitochondria were abundant in the pre-synaptic terminal (Figure 2).

**Figure 2.**
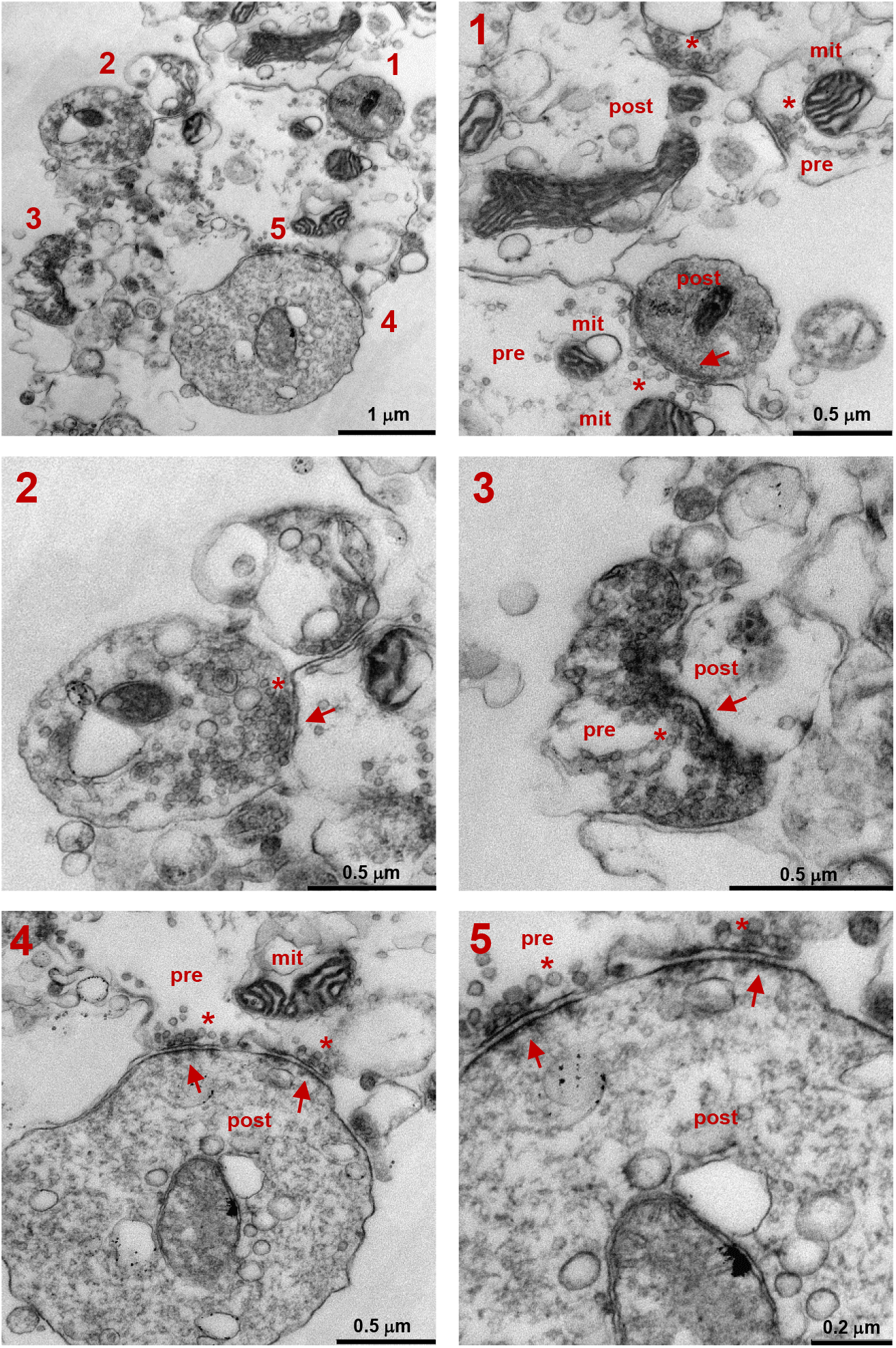
Electron micrographs of the 1P5 subcellular fraction. Pre-synaptic (pre) and post-synaptic (post) elements are marked, and the mitochondria (mit), synaptic vesicles (asterisk) and post-synaptic densities (arrows) can be distinguished.

### Characterization of the proteins present in SC SNs

The iTRAQ analysis carried out identified 3,133 proteins in SC SNs with at least one unique peptide (see below), applying a FDR = 0.01 (Table S2 in Supplemental File 1). Of these, 2,530 proteins were mapped to IDs in the PANTHER gene ontology tool. We carried out overrepresentation tests to classify these proteins according to different criteria: molecular function, protein class, cell component, pathways and biological processes in which these proteins are involved (Supplemental File 2). There was a significant representation of proteins from synapses, the cytoskeleton and the ATP synthase complex in the SC SNs (Figure 3). As such, SC SNs were enriched in proteins involved in the tricarboxylic acid (TCA) cycle and other oxidative metabolic processes, in cytoskeletal organizers and in proteins involved in synaptic signaling (for example, through metabotropic and ionotropic glutamate receptors, or dopamine, serotonin and acetylcholine receptors), as well as in proteins with other functions (Figure 3 and Supplemental File 2). Overall, these results corroborated the 1P5 fraction as the biochemical fraction enriched in SNs.

**Figure 3.**
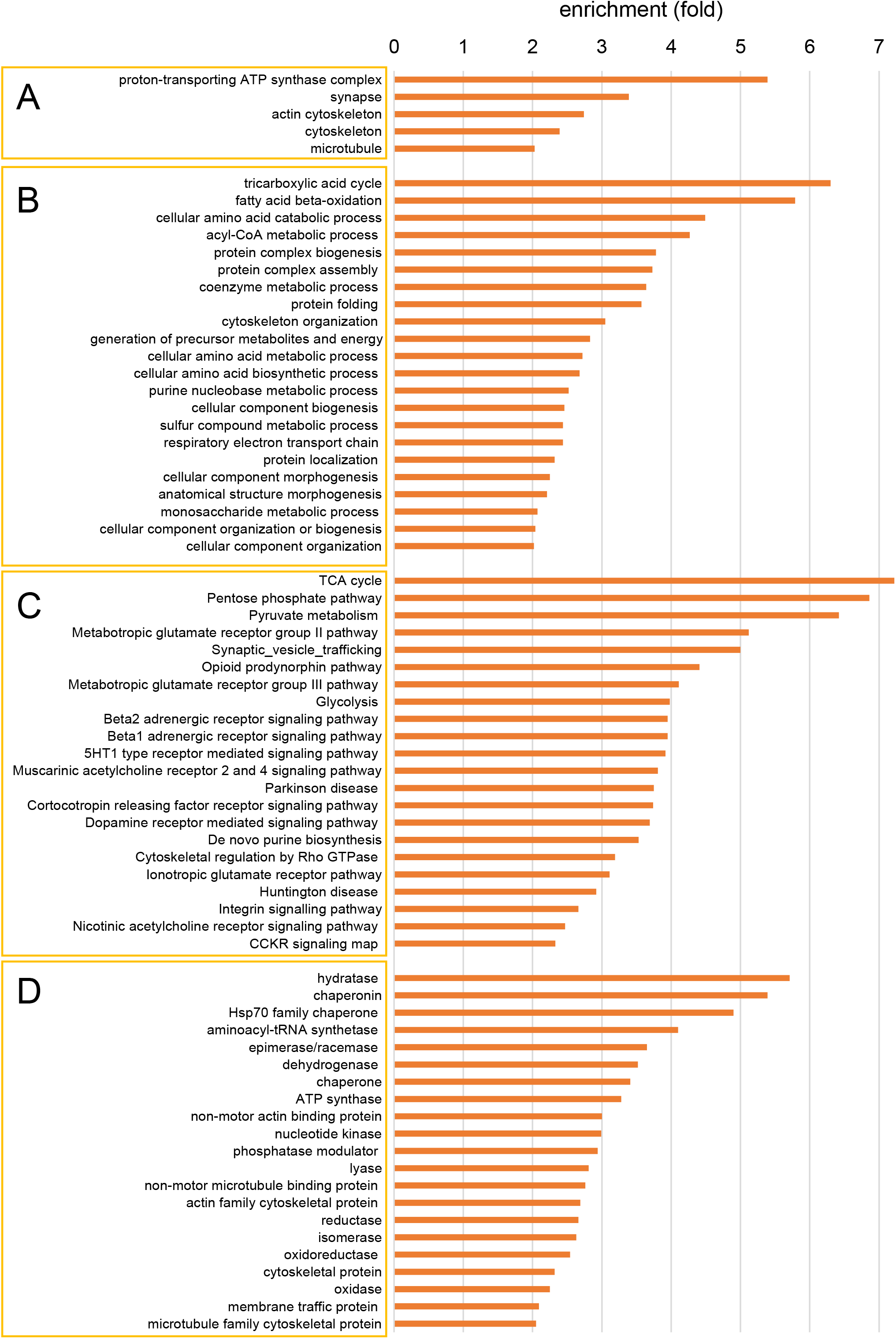
PANTHER overrepresentation tests of proteins detected in SC SNs by iTRAQ. The cellular components (A), biological processes (B), pathways (C) and protein classes (D) enriched more than 2-fold in SC SNs are shown.

Interestingly, protein folding was among the list of outstanding biological processes, and chaperones, aminoacyl-tRNA synthetases and RNA-binding proteins were among the protein classes enriched in SC SNs (Figure 3 and Supplemental File 2). Hence, mRNA transport, local synthesis and protein folding may be especially relevant for SC synaptic physiology.

### Relative protein levels in SC SNs from Tg-SOD1 mice, and Tg-SOD1/G93A mice at disease onset and symptomatic

To characterize the progress of SC synaptic failure in the ALS mouse model Tg-SOD1/G93A, we compared the relative levels of SN proteins at disease onset and in symptomatic Tg-SOD1/G93A animals. We isolated SC SNs from WT mice (littermates of Tg-SOD1/G93A animals that did not bear the transgene), from Tg-SOD1/G93A-ONSET mice in which the first signs of the disease have just appeared, and from overtly symptomatic Tg-SOD1/G93A-SYMPTOM animals. To control the effects associated to human SOD1 overexpression, SC SNs from Tg-SOD1 mice were also studied. These preparations were analyzed by iTRAQ proteomics and the amount of each protein relative to the WT SNs was defined (Table S2 in Supplemental File 1). Only proteins identified with at least two unique peptides were considered for quantitative analysis with the IPA software (1,652 proteins: Table S3 in Supplemental File 1), and a 2-fold cut-off was established to identify the most prominent protein changes. Using this criterion, only 5 proteins were found to be strongly deregulated in Tg-SOD1 SNs (3 up- and 2 down-regulated: Table S4 in Supplemental File 3). In the Tg-SOD1/G93A-ONSET sample, 69 proteins were affected (2 up- and 67 down-regulated: Table S5 in Supplemental File 3), while in the SNs from the Tg-SOD1/G93A-SYMPTOM mice the levels of most of these proteins (1,387 proteins) was altered (3 up- and 1,384 down-regulated: Table S6 in Supplemental File 3). To confirm these results, we examined the amount of the glutamate ionotropic receptor AMPA type 1 subunit (GRIA1) and dystroglycan 1 (DAG1) in Western blots of WT and Tg-SOD1/G93A-ONSET SNs. As expected, the levels of these proteins were reduced in Tg-SOD1/G93A-ONSET SNs (not shown).

While no relevant network of altered proteins was found in Tg-SOD1 SNs, two networks gave high scores in Tg-SOD1/G93A-ONSET SNs. Of these, Network 1 (score = 27) mainly included proteins involved in cell death and RNA expression (Figure 4A), whereas the proteins in Network 2 (score = 25) essentially participate in defining the morphology of neurons, movement disorders and in neuromuscular disease (Figure 4B). Interestingly, the IPA revealed that MNK1 was the most statistically significant upstream regulator (Z-score = −2.449, p value of overlap = 7.5E-06), suggesting that MNK1 downregulation could explain many of the protein alterations observed in Tg-SOD1/G93A-ONSET SC SNs.

**Figure 4.**
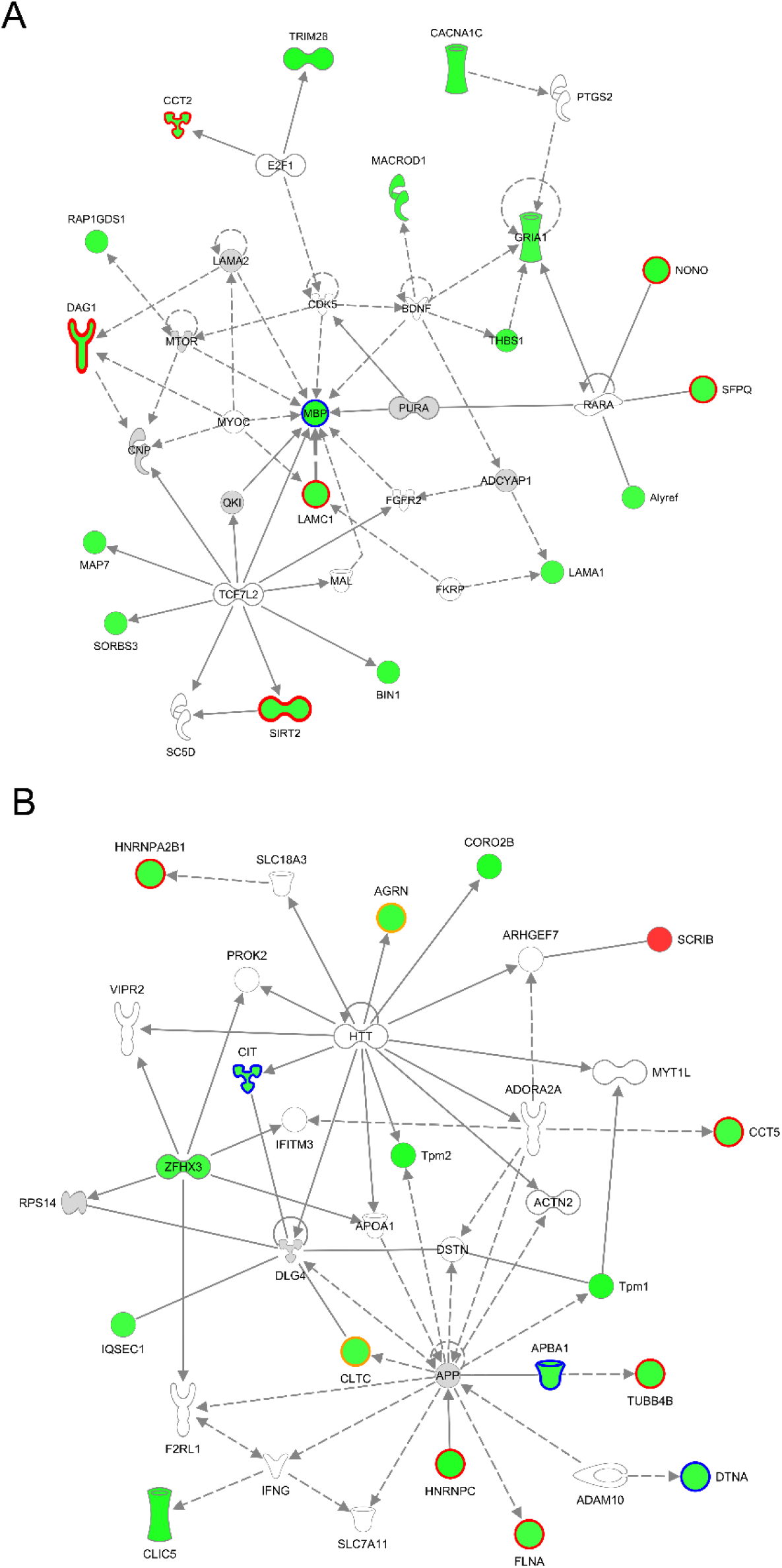
Networks of proteins affected in Tg-SOD1/G93A-ONSET SC SNs. (A) Network 1 (score = 27) is comprised of 35 proteins, of which 18 were downregulated (green) in Tg-SOD1/G93A-ONSET SNs. The proteins in grey were also detected in Tg-SOD1/G93A-ONSET SC SNs but they were unaffected considering a 2-fold cut-off. Continuous lines represent direct relationships between proteins and the discontinuous lines indicate indirect relationships. Proteins outlined in red are MNK1 targets, those outlined in blue are regulated by FMRP. In this network, 17 proteins are known to be involved in RNA expression, including MBP, NONO, SFPQ, SIRT2, SORBS3 and TRIM28. In addition, 26 proteins of this network are known to be involved in cell death, including BIN1, CACNA1C, CCT2, DAG1, LAMA1, MBP, RAP1GDS1, SFPQ, SIRT2, THBS1 and TRIM28. (B) Network 2 (score = 25) is comprised of 35 proteins, of which 16 were downregulated (green) and 1 was upregulated (red) in Tg-SOD1/G93A-ONSET SNs. The proteins in grey were also detected in Tg-SOD1/G93A-ONSET SC SNs but they were unaffected considering a 2-fold cut-off. Continuous lines represent direct relationships between proteins and discontinuous lines indicate indirect relationships. The proteins outlined in red are MNK1 targets, those outlined in blue are regulated by FMRP, and the proteins outlined in orange are regulated by MNK1 and FMRP. There were 14 proteins in this network known to be involved in movement disorders, including AGRN, APBA1, CIT, CORO2B, DTNA and TUBB4B. Moreover, 12 proteins in this network are known to be involved in neuronal morphology, including AGRN, APBA1, CLTC, DTNA, FLNA and HNRNPA2B1.

MNK1 is a kinase in the Ras-ERK signaling pathway that is involved in local translation and it mediates the BDNF-induced release of CYFIP1 from eIF4E in cortical neurons, triggering protein synthesis [29]. Indeed, these authors found that the synthesis of 718 proteins appeared to be regulated by MNK1 and, given that CYFIP1 interacts with FMRP [30], a significant overlap between these MNK1 regulated proteins and 842 high-confidence FMRP targets [31] was detected [29]. Thus, we compared the complete list of MNK1-regulated proteins [29] and the list of high-confidence FMRP-regulated proteins [31] with the list of proteins deregulated in Tg-SOD1/G93A-ONSET SC SNs. Remarkably, 22 proteins affected in Tg-SOD1/G93A-ONSET SC SNs (i.e.: 31.9%) were identified as MNK1 targets, and 8 proteins (i.e.: 11.6%) were identified as FMRP targets (Table 1). These data suggest a possible role of MNK1 (and/or FMRP) in the onset of ALS.

**Table 1.**
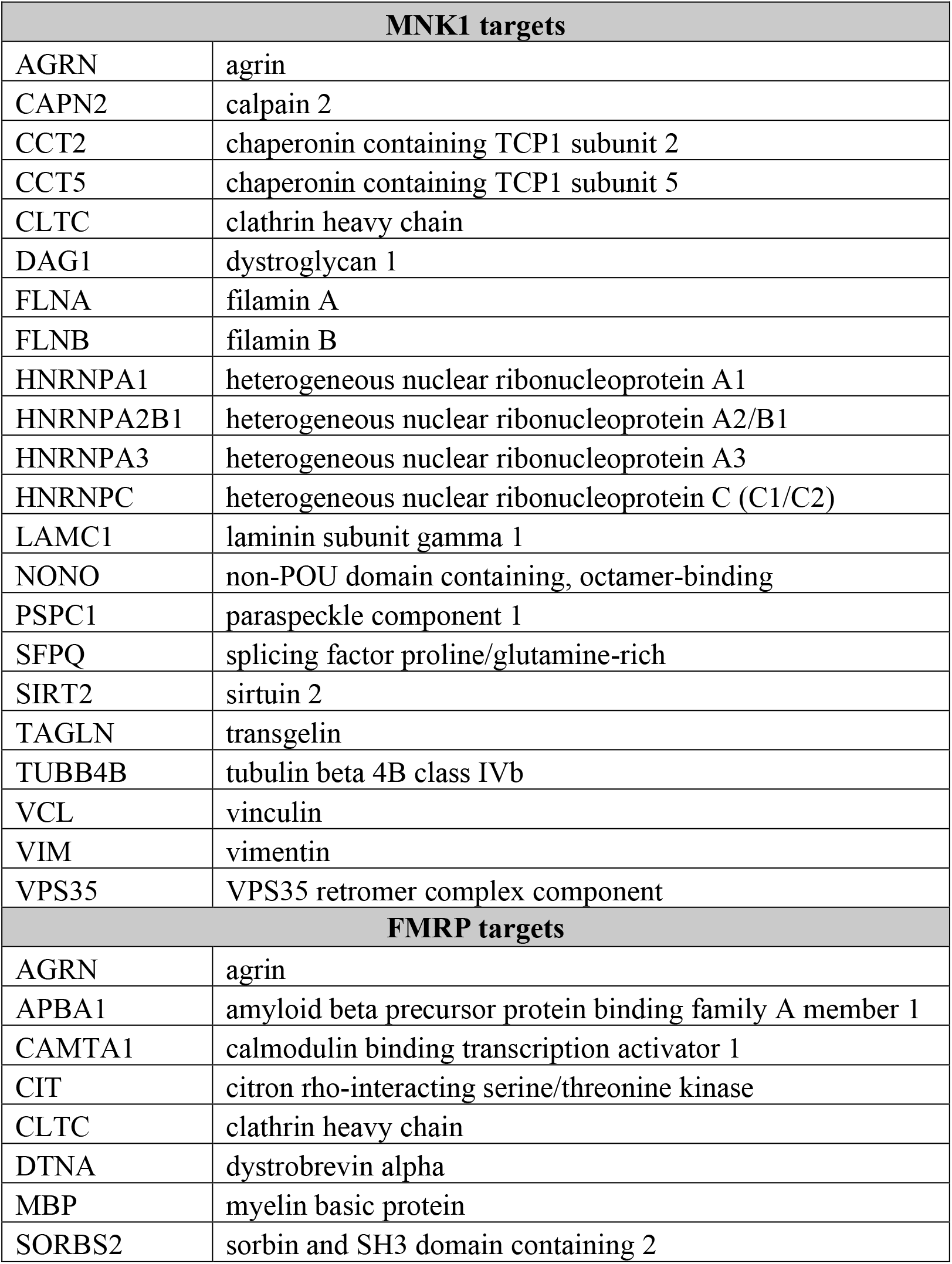
Proteins altered in G93A-ONSET SC SNs known to be regulated by MNK1 or FMRP.

| **MNK1 targets** |  |
| --- | --- |
| AGRN | agrin |
| CAPN2 | calpain 2 |
| CCT2 | chaperonin containing TCP1 subunit 2 |
| CCT5 | chaperonin containing TCP1 subunit 5 |
| CLTC | clathrin heavy chain |
| DAG1 | dystroglycan 1 |
| FLNA | filamin A |
| FLNB | filamin B |
| HNRNPA1 | heterogeneous nuclear ribonucleoprotein A1 |
| HNRNPA2B1 | heterogeneous nuclear ribonucleoprotein A2/B1 |
| HNRNPA3 | heterogeneous nuclear ribonucleoprotein A3 |
| HNRNPC | heterogeneous nuclear ribonucleoprotein C (C1/C2) |
| LAMC1 | laminin subunit gamma 1 |
| NONO | non-POU domain containing, octamer-binding |
| PSPC1 | paraspeckle component 1 |
| SFPQ | splicing factor proline/glutamine-rich |
| SIRT2 | sirtuin 2 |
| TAGLN | transgelin |
| TUBB4B | tubulin beta 4B class IVb |
| VCL | vinculin |
| VIM | vimentin |
| VPS35 | VPS35 retromer complex component |
| **FMRP targets** |  |
| AGRN | agrin |
| APBA1 | amyloid beta precursor protein binding family A member 1 |
| CAMTA1 | calmodulin binding transcription activator 1 |
| CIT | citron rho-interacting serine/threonine kinase |
| CLTC | clathrin heavy chain |
| DTNA | dystrobrevin alpha |
| MBP | myelin basic protein |
| SORBS2 | sorbin and SH3 domain containing 2 |

As most proteins were affected in the Tg-SOD1/G93A-SYMPTOM SC SNs, it would appear that there was a general perturbation of synaptic proteins, probably related to the motor neurodegeneration that is characteristic of advanced ALS stages. Accordingly, IPA predicted an impairment in synaptic transmission and in the transport of synaptic vesicles, as well as enhanced neurodegeneration in these mice (not shown).

### A reduction in the MNK1 protein in SC SNs from Tg-SOD1/G93A mice at the onset of the disease

The IPA analysis suggested that reduced levels and/or activity of MNK1 could be responsible for the proteomic changes in Tg-SOD1/G93A-ONSET SC SNs. As the MNK1 protein was not detected in the iTRAQ experiments, we quantified the amount of MNK1 protein in Western blots of SNs from WT and Tg-SOD1/G93A-ONSET mice. As predicted by the IPA, reduced MNK1 levels were detected in the Tg-SOD1/G93A-ONSET SNs relative to the WT SNs (Figure 5A). This reduction was specific to the synaptic preparations as it was not observed in the total protein extracts (Figure 5B).

**Figure 5.**
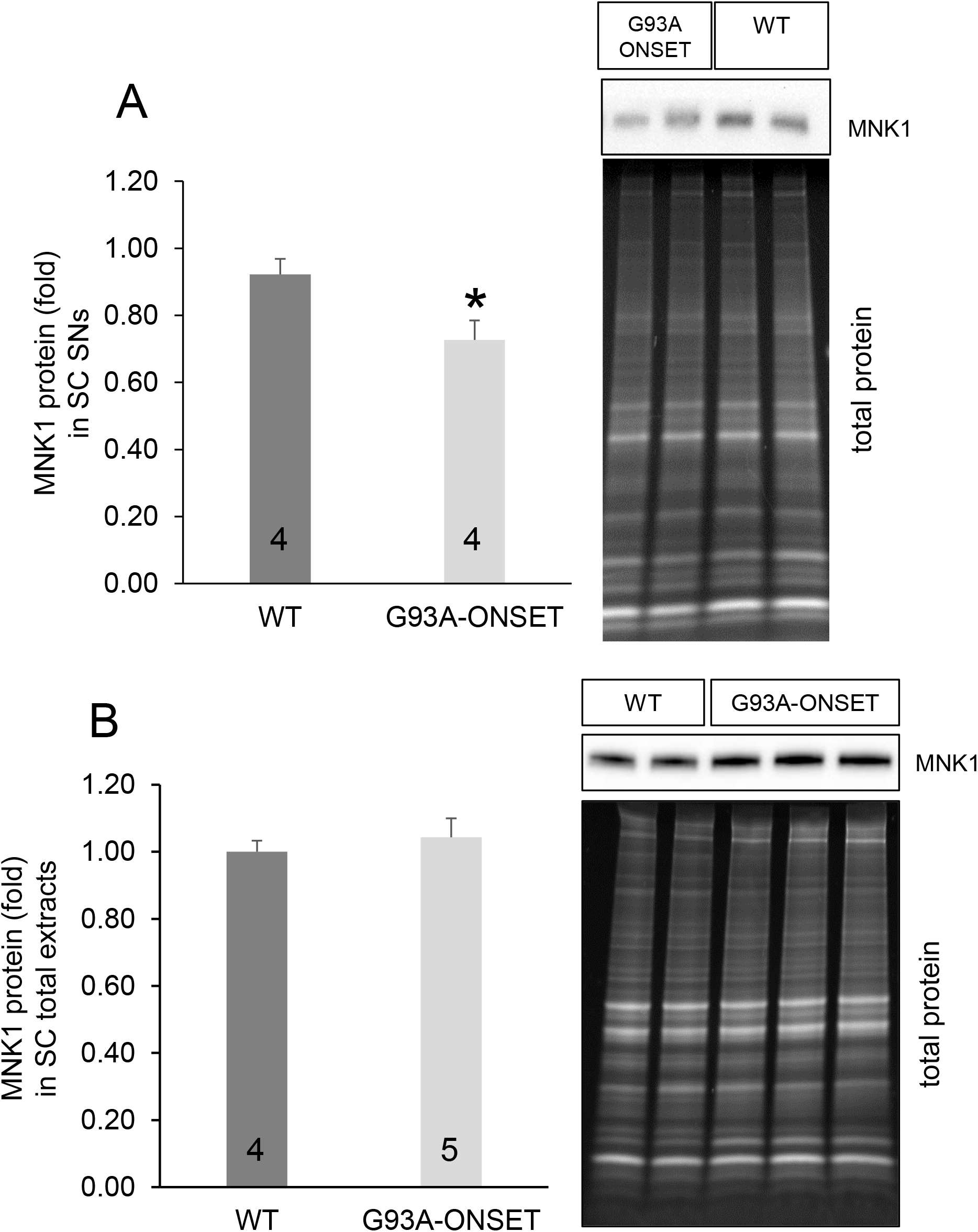
Levels of MNK1 protein are reduced in SC SNs but not in the total protein extracts of Tg-SOD1/G93A-ONSET mice. (A) Proteins from WT and Tg-SOD1/G93A-ONSET SC SNs were analyzed in Western blots probed with an anti-MNK1 antibody. The signals obtained were normalized to the total protein loaded and the mean values are shown, with the number of independent SN preparations indicated in the corresponding bars. The error bars indicate the SEM and the asterisk denotes a statistically significant difference (p=0.039, t-test). A representative Western blot is shown. (B) Proteins from WT and Tg-SOD1/G93A-ONSET SC total extracts were analyzed in Western blots probed with an anti-MNK1 antibody. The signals obtained were normalized to the total protein loaded and the mean values are shown, with the number of independent total protein extracts analyzed indicated in the corresponding bar. The error bars indicate the SEM and a representative Western blot is shown.

### Lower rates of translation in SC SNs from Tg-SOD1/G93A-ONSET mice

Since MNK1 is involved in local synaptic translation, we analyzed translation in SC SNs from Tg-SOD1/G93A mice. As such, metabolic labeling revealed an average 30% reduction in the translation rate in G93A-ONSET SC SNs (mean translation rate 69.6 ± 4.5%, n=7) relative to WT SC SNs (Figure 6). A similar reduction in metabolic labeling was also evident in SC SNs from symptomatic mice (mean translation rate 66.0 ± 15%, n=2), whereas normal or even enhanced labeling was seen in Tg-SOD1 SC SNs (Figure 6). In conclusion, these data suggest that local synaptic translation is impaired in the Tg-SOD1/G93A SC at early stages of the disease.

**Figure 6.**
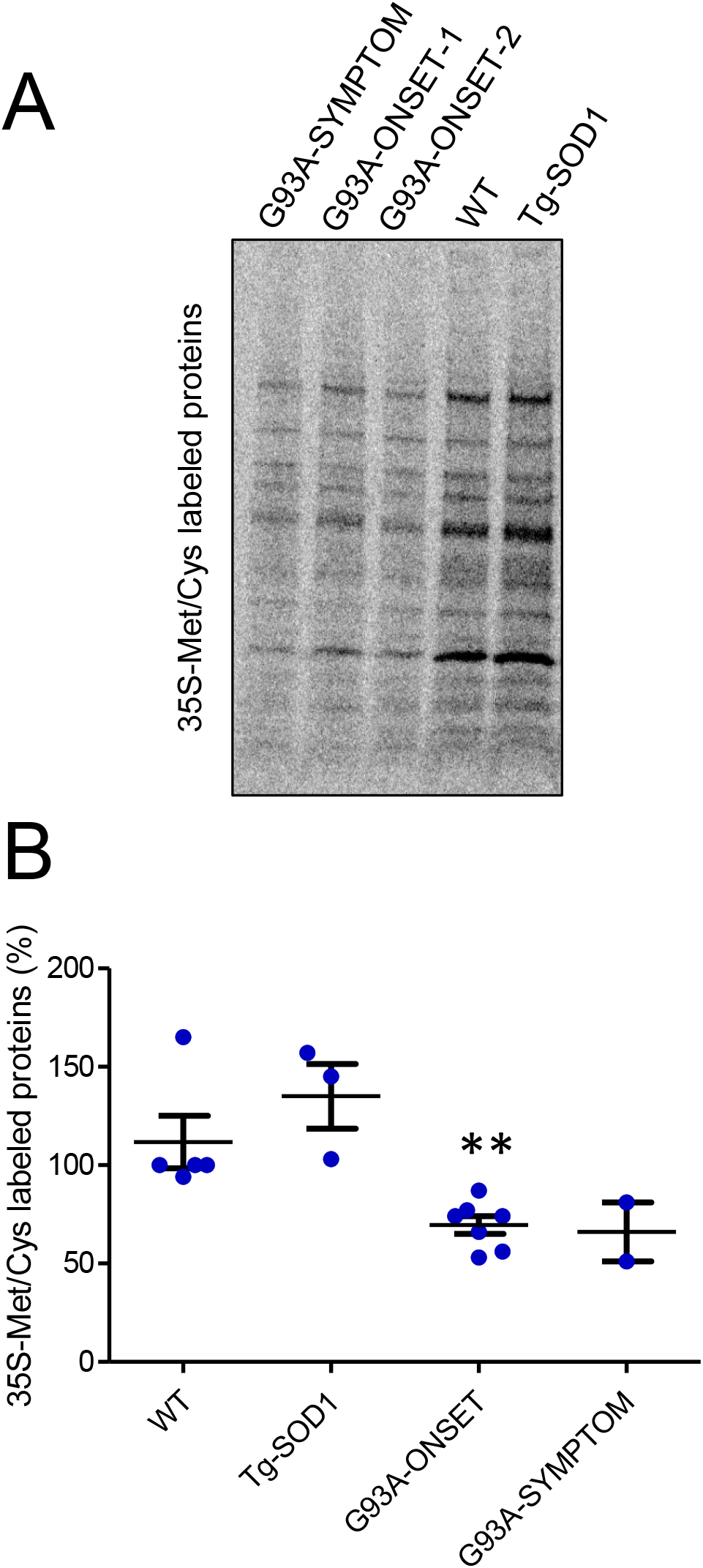
Local synaptic translation is reduced in the SC of Tg-SOD1/G93A-ONSET mice. (A) ^35^S-Met/Cys radioactive labeled proteins in the SC SNs from symptomatic Tg-SOD1/G93A-SYMPTOM mice, Tg-SOD1/G93A-ONSET mice at disease onset (ONSET 1 and 2), and from wild type (WT) and Tg-SOD1 mice. Similar amounts of protein were loaded for each condition and a representative experiment is shown. (B) Quantification of the radioactive, neo-synthesized proteins in WT, Tg-SOD1 and Tg-SOD1/G93A SC SNs from three independent experiments. The radioactivity incorporated into neo-synthesized proteins was quantified in SC SNs obtained from WT (n=5), Tg-SOD1 (n=3) and Tg-SOD1/G93A-ONSET (n=7) or Tg-SOD1/G93A-SYMPTOM (n=2) mice, normalized to the total protein content of the corresponding sample. The data are shown as the percentages compared to the WT values and the asterisks indicate significant difference between Tg-SOD1/G93A-ONSET and WT SNs (p=0.0065, t-test).

## DISCUSSION

Since their initial description [32], SNs have been used extensively to explore different aspects of synaptic physiology. Indeed, more than 14,000 publications can be retrieved from PubMed by searching with the word “synaptoneurosome”. Generic protocols are often used to isolate SNs regardless of the specie (mouse, rat or other) or the nervous system region studied, two factors that will critically affect the outcome. Here we show that, using the protocol we originally established to isolate SNs from the mouse hippocampus and cortex, SC SNs (as characterized biochemically and morphologically) localize to a different gradient fraction.

Through iTRAQ proteomics, we found that mitochondrial, synaptic and cytoskeletal proteins were among the most strongly represented proteins in SC SNs, as previously reported in other proteomics studies focused on synaptic cortical preparations (for example see [33, 34]). Interestingly, we found that chaperonins and chaperones, especially those belonging to the Hsp70 family, were particularly overrepresented in SC SNs. Remarkably, Hsp70 chaperones (and their Hsp40 partners) fulfil an important role in translation, associating with polysomes and enhancing translation by promoting the folding of nascent peptides (for a review see [35]). In relation to this, aminoacyl-tRNA synthetases were also overrepresented in SC SNs, which could reflect the relevant role of these enzymes in translation at SC synapses, as is indeed the case in hippocampus [34]. Hence, it is tempting to speculate that local translation could be particularly important for motor neurons and consequently, motor neurons could be particularly sensitive to any defects in local translation.

Since ALS is diagnosed when the first clinical manifestations appear, we were interested in determining the proteins that experimented alterations early in the disease. Thus, we focused on Tg SOD1/G93A mice at the onset of the symptoms. Strikingly, of the 69 proteins altered in SC SNs from Tg-SOD1/G93A-ONSET mice, many of them were involved in the processing or binding of mRNA and/or translation. In fact, we found that a significant percentage of the affected proteins were targets of MNK1, a key regulator of local dendritic translation. MNK1 was predicted by IPA to be the most significant upstream regulator of the set of proteins altered in Tg-SOD1/G93A-ONSET SC SNs and it was predicted to be downregulated in the SC SNs from these mice. Accordingly, we found less MNK1 protein in SC SNs and reduced local translation rates (measured as the radioactive labelling of neo-synthetized proteins in SNs). Among the proteins identified, several were of particular interest in the context of ALS. Thus, mutations in the mRNA binding proteins HnRNPA1, HnRNNPA2/B1 or HnRNPA3 seem to cause ALS [23, 36). In addition, the retromer complex subunit VPS35 has recently been seen to be reduced in motor neurons of asymptomatic and symptomatic G93A mice, and in ALS patients. Indeed, retromer stabilization by increasing VPS35 levels ameliorates the phenotype of G93A mice [37].

It is well known that, in response to BDNF or glutamate, local translation is activated through two related pathways: the Akt-mTOR and the Ras-ERK pathways (for a review see [38]). MNK1 functions in the Ras-ERK branch, phosphorylating eIF4E and thereby triggering translation. In addition, MNK1 activation disrupts the inhibitory complex formed by CYFIP1 and eIF4E, triggering the translation of specific targets [29]. Thus, in basal conditions FMRP tethers target mRNAs to CYFIP1, which blocks eIF4E and prevents the initiation of translation [30]. TrkB or mGluR activation provokes the MNK1-dependent release of CYFIP1 from eIF4E, inducing the translation of some FMRP targets [29, 30]. Therefore, the reduced synaptic levels of MNK1 detected in Tg-SOD1/G93A-ONSET mice may not only affect general local translation but more specifically, FMRP-regulated local translation. As proposed previously [29], the complex molecular relationships described above could explain the significant overlap between MNK1 and FMRP targets. Indeed, we found that a number of FMRP targets were also affected in SC SNs from Tg-SOD1/G93A-ONSET mice (Table 1). As mentioned, FMRP acts primarily as a translational repressor but it has also been demonstrated to directly activate SOD1 translation by binding to a particular structural motif located in the SOD1 5’UTR, the SoSLIP motif [15]. Given the molecular relationships between MNK1 and FMRP [29], and between FMRP and SOD1 [15], it is tempting to speculate that aberrant dendritic SOD1 translational complexes in Tg-SOD1/G93A-ONSET mice might sequester FMRP (and perhaps chaperones), which could interfere with translation of other FMRP target mRNAs, MNK1 targets and/or with local translation in general. In ALS mouse models, aberrant SOD1 aggregates first appear in dendrites [9–11] and dendritic aggregates have also been described in ALS patients [12, 13]. Interestingly, it has been reported that WT and mutant SOD1/G93A proteins localize to stress granules and P-bodies, and that SOD1/G93A inclusions also contain mRNAs [39]. Although the precise molecular mechanisms responsible for the reduced synaptic levels of MNK1 remain to be elucidated, our findings point to an important role of MNK1 and local translation in the early stages of ALS. Thus, MNK1 could represent a potential therapeutic target in ALS.

## ACKNOWLEDGMENTS

This work was supported by the Junta de Andalucía (Grant P09-CTS-4610). We thank Juan Luis Ribas (CITIUS, Microscopy Service, Universidad de Sevilla, Spain) and Mariló Pastor (IBiS Proteomics Service, Instituto de Biomedicina de Sevilla, Spain) for their technical help and advice. We also thank the Supercomputing and Bioinnovation Center (Universidad de Málaga, Spain) for providing us with access to the IPA tool.

## SUPPLEMENTAL DATA

**Supplemental File 1:**

**- Supplemental Table S1.** Criteria based on the phenotypic traits used to classify the Tg-SOD1/G93A mice as at disease onset or symptomatic.
**- Supplemental Table S2.** iTRAQ proteomics - raw data.
**- Supplemental Table S3.** iTRAQ proteomics data after removing proteins identified with only one unique peptide.

**Supplemental File 2:** PANTHER overrepresentation tests for cellular components, biological processes, pathways, protein classes and molecular functions of proteins identified by iTRAQ in SC SNs.

**Supplemental File 3:**

**- Supplemental Table S4.** Proteins altered in Tg-SOD1 SC SNs.
**- Supplemental Table S5.** Proteins altered in Tg-SOD1/G93A-ONSET SC SNs.
**- Supplemental Table S6.** Proteins altered in Tg-SOD1/G93A-SYMPTOM SC SNs.

